# NF-κB-Inducing Kinase (NIK) Governs the Mitochondrial Respiratory Capacity, Differentiation, and Inflammatory Status of Innate Immune Cells

**DOI:** 10.1101/2022.08.06.503047

**Authors:** Justin N. Keeney, Ashley Winters, Raquel Sitcheran, A. Phillip West

## Abstract

NF-κB-Inducing Kinase (NIK), which is essential for the activation of the noncanonical NF-κB pathway, regulates diverse processes in immunity, development, and disease. While recent studies have elucidated important functions of NIK in adaptive immune cells and cancer cell metabolism, the role of NIK in metabolic-driven inflammatory responses in innate immune cells remains unclear. Here, we demonstrate that NIK-deficient bone marrow-derived macrophages exhibit defects in mitochondrial-dependent metabolism and oxidative phosphorylation (OXPHOS), which impairs the acquisition of a pro-repair, anti-inflammatory phenotype. Subsequently, NIK-deficient mice exhibit skewing of myeloid cells characterized by aberrant eosinophil, monocyte, and macrophage cell populations in the blood, bone marrow, and adipose tissue. Furthermore, NIK-deficient blood monocytes display hyperresponsiveness to bacterial lipopolysaccharide and elevated TNFα production ex vivo. These findings suggest that NIK governs metabolic rewiring, which is critical for balancing pro- and anti-inflammatory myeloid immune cell function. Overall, our work highlights a previously unrecognized role for NIK as a molecular rheostat that fine-tunes immunometabolism in innate immunity and suggests that metabolic dysfunction may be an important driver of inflammatory diseases caused by aberrant NIK expression or activity.

**Key Points:**

- NIK-deficient macrophages exhibit impaired mitochondrial oxidative phosphorylation
- NIK-deficient mice have more inflammatory myeloid cells in blood and bone marrow
- NIK-dependent metabolic rewiring shapes pro- and anti-inflammatory innate immunity

## Introduction

Macrophages are important components of the innate immune system that play key roles in immunity, tissue homeostasis, and cancer (1). Macrophages exist across a spectrum of polarization states in vivo and can adopt pro-inflammatory or anti-inflammatory properties depending on local cytokine gradients and tissue residency (2). Murine bone marrow-derived macrophages (BMDMs) can be polarized toward the extreme end of the pro-inflammatory spectrum by culture in bacterial lipopolysaccharide (LPS) or interferon gamma. These so-called M1 or M(LPS) macrophages produce pro-inflammatory cytokines such as TNFα, have bactericidal and phagocytic functionality, and can produce reactive oxygen species (ROS) and nitric oxide (NO) (3). Conversely, BMDMs cultured in interleukin-4 (IL-4) are skewed toward the anti-inflammatory end of the spectrum. These so-called M2 or M(IL-4) macrophages can dampen inflammatory responses by secreting anti-inflammatory cytokines such as interleukin-10 (IL-10), promote extracellular matrix remodeling, and enhance stromal cell proliferation (3). It is well appreciated that metabolic switches promote the acquisition of distinct macrophage polarization states and play pivotal roles in the final effector cell functionality (4). M1 macrophages upregulate glycolysis, have a broken tricarboxylic acid (TCA) cycle, and display impaired oxidative phosphorylation (OXPHOS), leading to enhanced generation of TCA intermediates such as succinate and other metabolites necessary for ROS and NO generation (5– 7). On the other hand, M2 macrophages have an intact TCA cycle and upregulate OXPHOS and fatty acid oxidation to generate anti-inflammatory products such as glucocorticoids, IL-10, IL-13, etc. (8, 9). These metabolic shifts are critical for macrophage function, as small molecules that inhibit the major metabolic enzymes in glycolysis, TCA, and OXPHOS can dramatically alter polarization states in vitro and in vivo (10–12).

Nuclear factor-kappaB (NF-κB) signaling has a well-established role in inflammatory processes with the canonical NF-κB pathway rapidly responding to inflammatory stimuli (13). Alternatively the non-canonical NF-κB pathway, which requires NF-κB Inducing Kinase (NIK; also known as MAP3K14), regulates immune cell differentiation, development, and tissue homeostasis (13–16). Over the last decade, NIK has been found to play many novel roles in a diverse set of immune cells and their progenitors (17, 18). Within the bone marrow, NIK promotes mitochondrial biogenesis during osteoclastogenesis (14). In microglia, which are tissue-resident macrophages of the brain, NIK is important for the induction of cytokine expression, which aids in T-cell recruitment in an animal model of experimental autoimmune encephalomyelitis (19, 20).

Recently, we and others have demonstrated roles for NIK in regulating mitochondrial function and metabolism in cancer cells (19, 21–23). For example, we previously reported that NIK regulates mitochondrial dynamics (21) and promotes oxidative metabolism (22) in glioblastoma, an aggressive form of brain cancer. Moreover, NIK-deficient glioblastoma cells exhibit impaired mitochondrial fission and increased glycolysis to compensate for diminished oxidative metabolism (21, 22) and are less invasive. Notably, we have shown that these novel metabolic roles of NIK are largely independent of non-canonical NF-κB signaling, suggesting that NIK plays a direct role in regulating mitochondrial function. Another recent study showed that NIK shapes the metabolism of CD8 T cells independently of non-canonical NF-κB by stabilizing hexokinase 2 and promoting NADPH generation (23). Similar results were found in B cells containing a deletion of TRAF3, an upstream inhibitor of NIK (24). Although NIK has emerged as a key regulator of tumor cell and lymphocyte metabolism, whether NIK regulates immunometabolic switches in myeloid innate immune cells remains unclear.

Studies using NIK knockout (NIK^KO^) mice have revealed that NIK deficiency promotes a dramatic increase in peripheral blood eosinophil abundance (25). NIK^KO^ mice develop a hypereosinophilic inflammatory disease driven by the infiltration of pathogenic eosinophilic granulocytes into diverse organs, including skin, liver, lung, and esophagus, leading to early death (25, 26). Recent studies have shown that eosinophils upregulate OXPHOS upon activation (27); however, it remains unclear whether immunometabolic switches contribute to the inflammatory myeloid phenotypes observed in NIK^KO^ mice. To begin to address this knowledge gap, we measured mitochondrial and metabolic parameters in BMDMs from NIK^KO^ mice. Here, we show that NIK promotes oxidative metabolism in macrophages and that loss of NIK results in hyperinflammatory myeloid skewing of the bone marrow, blood, and other tissues. Our findings demonstrate that NIK is an important molecular rheostat that maintains mitochondrial function to control innate immune cell function, and they suggest that metabolic alterations may contribute to diseases where NIK activity is dysregulated.

## Materials and methods

### Mice

All animal experiments were performed in accordance with animal use protocols with approved IACUC guidelines at Texas A&M University. *Map3k14* null mice were purchased from The Jackson Laboratory (strain 025557) and maintained by heterozygous breeding. *Map3k14* floxed mice were obtained from Genentech and bred with LysMCre mice (Jackson Lab strain 004781). Equivalent numbers of male and female mice were used for all experiments. Animal numbers used are described in figure legends.

### Reagents

A comprehensive list of antibodies, reagents, and primers is provided in Tables 1-3, respectively.

**Table 1:** Primary antibodies and antibody-based reagents used in the study.

| Catalog Number | Company | Reagent | Conjugate |
| --- | --- | --- | --- |
| **Stemcell Panel** |  |  |  |
| A15423 | Invitrogen | Anti-MO CD117/C-Kit | APC-Cy7 |
| 15-5981-81 | Invitrogen | Anti-MO Ly-6A/E (SCA1) | PE-Cyn5 |
| 25-0481-80 | Invitrogen | Anti-MO CD48 | PE-Cyn7 |
| 12-1351-81 | Invitrogen | Anti-MO CD135 | PE |
| 17-1502-80 | Invitrogen | Anti-MO CD150 | APC |
| 11-0341-82 | Invitrogen | Anti-MO CD34 | FITC |
| **Immune Panel** |  |  |  |
| 25-0032-80 | Invitrogen | Anti-MO CD3 | PE-Cyn7 |
| 12-0452-81 | Invitrogen | Anti-MO CD45R(B220) | PE |
| 11-5931-81 | Invitrogen | Anti-MO Ly-6G | FITC |
| 35-0451-80 | Invitrogen | Anti-MO CD45 | PE-Cyn5.5 |
| 17-0112-81 | Invitrogen | Anti-MO CD11b | APC |
| 155505 | Biolegend | Anti-MO Siglec F | PE |
| **In vivo Polarization** |  |  |  |
| 117351 | Biolegend | Anti-MO CD11c | APC |
| 144509 | Biolegend | Anti-MO CD193 (CCR3) | FITC |
| 123111 | Biolegend | Anti-MO F4/80 | PE-Cyn5 |
| **Blood and Spleen Immune Phenotyping** |  |  |  |
| 25-0112-U100 | Tonbo Biosciences | Anti-HU/MO M1/70 (CD11b) | APC-Cy7 |
| 115539 | Biolegend | Anti-MO CD19 | BV605 |
| 100267 | Biolegend | Anti-MO CD3 | APC-Fire810 |
| 65-4801-U100 | Tonbo Biosciences | Anti-MO F4/80 | PerCP-C5.5 |
| 128049 | Biolegend | Anti-MO Ly6C | BV650 |
| 50-1276-U100 | Tonbo Biosciences | Anti-MO Ly-6G | PE |
| 107643 | Biolegend | Anti-MO MHCII | BV711 |
| 506324 | Biolegend | Anti-MO TNFα | PE-Cy7 |
| 13-0871-T100 | Tonbo Biosciences | Viability | Ghost Dye 710 |
| 70-0161-U100 | Tonbo Biosciences | Anti-MO CD16 / CD32 (Fc Shield) (2.4G2) |  |
| **ELISA Kit** |  |  |  |
| 41-9234-P002 | Tonbo Biosciences | Mouse IL-10 ELISA Matched Antibody Pair Kit |  |

| **Immunofluorescence** |  |  |  |
| --- | --- | --- | --- |
| MABT166 | Millipore | Anti-TFAM |  |
| CBL186 | Millipore | Anti-DNA |  |
| N-20 | Santa Cruz<br>Biotechnology | Anti-HSP60 |  |
| **Western Blotting** |  |  |  |
| 4994 | Cell Signaling | Anti-NIK |  |
| 2697 | Cell Signaling | Anti-pIKKα/IKKβ S176/180 | |
| OP133 | EMD Millipore | Anti-IKKα | |
| 4882 | Cell Signaling | Anti-p100/p52 |  |
| SC-365062 | Santa Cruz<br>Biotechnology | Anti-GAPDH |  |
| SC-69879 | Santa Cruz<br>Biotechnology | Anti-β-Actin | |
| SC-5274 | Santa Cruz<br>Biotechnology | Anti-β-Tubulin | |
| 4661T | Cell Signaling | Anti-VDAC |  |

**Table 2:** Commercial reagents used in the study.

| **Catalog Number** | **Supplier** | **Reagent** |
| --- | --- | --- |
| L31963 | Invitrogen | Violet Live/Dead dye |
| 14-9161-71 | Invitrogen | HU FCR Binding Inhibitor |
| M36008 | Invitrogen | MitoSox |
| M22425 | Invitrogen | Mitotracker Red FM |
| M7514 | Invitrogen | Mitotracker Green FM |
| 8778s | Cell Signaling | Mitotracker Deep Red FM |
| TLRL-PB5LPS | Invivogen | Ultrapure LPS-B5 |
| 214-14 | PeptoTech | Recombinant Murine IL-4 |
| LS004174 | Worthington Biochemical | Collagenase, Type 2 |
| LS004186 | Worthington Biochemical | Collagenase, Type 4 |
| 354234 | Corning | Matrigel Basement Membrane Matrix |
| F3879 | Sigma Aldrich | Fibrinogen |
| 341584 | Sigma Aldrich | Polyvinyl Alcohol |
| C35007 | Invitrogen | CyQUANT |
| D275 | Invitrogen | DiO |
| F3506 | Sigma Aldrich | FMLP |
| TNB-0607-KIT | Tonbo Biosciences | Foxp3/Transcription Factor Staining Buffer Kit |
| 75351 | Sigma Aldrich | Oligomycin A |
| 15218 | Cayman Chemical | FCCP |
| A8674 | Sigma Aldrich | Antimycin A |
| ALX-350-360-G001 | Enzo Life Sciences | Rotenone |
| 1708891 | BioRad | iScript |
| A25742 | Universal SYBR Green | Invitrogen |
| 700930 | Cayman Chemical | MitoCheck Complex I Activity Assay Kit |
| 700940 | Cayman Chemical | MitoCheck Complex II Activity Assay Kit |
| 700950 | Cayman Chemical | MitoCheck Complex II/III Activity Assay Kit |
| 700990 | Cayman Chemical | MitoCheck Complex IV Activity Assay Kit |
| 701000 | Cayman Chemical | MitoCheck Complex V Activity Assay Kit |
| 5120-0075 | KPL | TMB Substrate |
| 15702-10 | Polysciences Inc | 1um carboxylate microspheres - FITC |
| HY-120501 | MedChem Express | B022 NIK inhibitor |
| P7170-1 | Sigma Aldrich | Ponceau S |
| 24592 | Thermo Scientific | GelCode Blue |

**Table 3:**
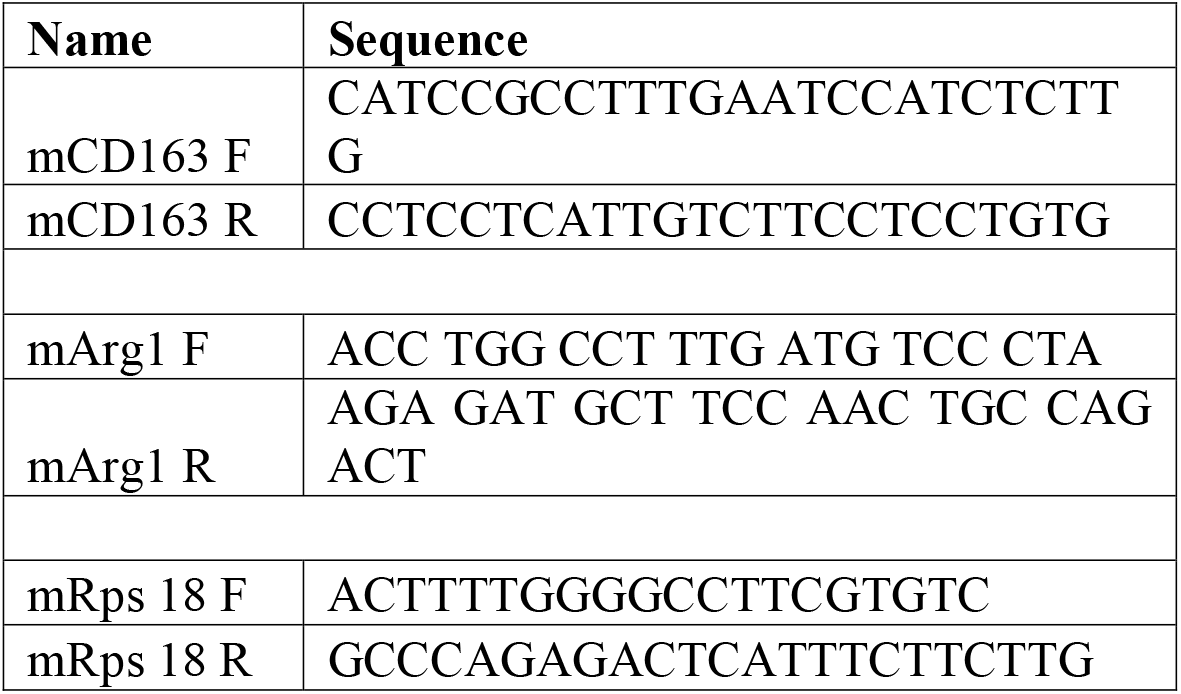
Oligonucleotides used in quantitative real-time PCR.

### Flow Cytometry

#### Blood and spleen immune phenotyping

Whole mouse blood was collected in sodium heparin tubes. Spleens were collected and filtered through a 100uM cell strainer. RBCs were lysed twice with ACK lysis buffer, and leukocytes were subjected to LPS stimulation (1 μg/ml) in the presence of brefeldin A and monensin for 4 hours. Fc receptors were blocked with anti-mouse CD16/CD32 (2.4G2) antibody, and cells were stained with antibodies against surface proteins, permeabilized with Foxp3/Transcription Factor Staining Buffer Kit (TNB-0607-KIT, Tonbo Biosciences, San Diego, CA, USA), and stained with antibodies against intracellular proteins. Cells were analyzed with a Cytek Aurora 5L Spectral Analyzer. The results were plotted and analyzed using FlowJo software (BD Bioscience; Franklin Lakes, NJ, USA).

#### Bone marrow phenotyping

Cells were isolated from the tibia and femurs of mice as described below (see Cell Culture). After an ACK lysis step, 500,000 cells per mouse were taken for immune cell phenotyping and 1 million cells were stained for the stem cell analysis. Cells were stained in a 96 well plate with Live/Dead Fix Violet (eBioscience, Waltham, MA, USA) at 4°C for 20 minutes. After being washed thoroughly, cells were blocked with FC blocker (eBioscience, Waltham, MA, USA) for 10 minutes at 4°C before the addition of the primary antibody cocktail which incubated with the cells for 30 minutes. After, cells were washed and fixed with 4% paraformaldehyde for 10 minutes at room temperature in the dark. Flow cytometric analysis was performed on a BD Fortessa X-20 (BD Bioscience; Franklin Lakes, NJ, USA). The results were plotted and analyzed using FlowJo software (BD Bioscience; Franklin Lakes, NJ, USA).

#### Fat and small intestine immune phenotyping

Visceral fat deposits and a 5cm section of the ileum were harvested from mice and washed with PBS before being placed in a 0.2% (2mg/mL) collagenase 2 solution, (Worthington Biochemical; Lakewood, NJ, USA) diluted in RIPA buffer (Gibco, ThermoFisher Scientific; Waltham, MA, USA), under contestant stirring for 60 minutes at 37°C. The isolated cells were filtered through a 70 μM cell strainer and red blood cells were lysed with ACK lysis buffer (VWR; Radnor, PA, USA). Cells were then stained as described above for the bone marrow.

#### Mitochondrial flow cytometry assays

This assay was previously described (28), but in brief: NIK^WT^ and NIK^KO^ BMDMs were treated with 20ng/mL IL-4 for 5 hours before cells were detached using PBS containing 1mM EDTA (Gibco, ThermoFisher Scientific; Waltham, MA, USA). Cells were then stained with a combination of Mitotracker Green FM, Mitotracker Red FM, Mitosox, Anti-MO CD11b, and Violet live dead dye for 15 minutes in a 37°C tissue culture incubator. Cells were then washed twice before analysis on a BD Fortessa X-20 flow cytometer. The results were plotted and analyzed using FlowJo software (BD Bioscience; Franklin Lakes, NJ, USA).

### Cell Culture

L929 and HEK293T cells were obtained from the American Type Culture Collection (ATCC) and maintained in Dulbecco’s Modified Eagle’s Medium (DMEM; Gibco, ThermoFisher Scientific; Waltham, MA, USA) supplemented with 10% fetal bovine serum (FBS; VWR; Radnor, PA, USA). BMDMs were generated by crushing the tibia and femurs of unpooled mice. After lysis of red blood cells with ACK lysis buffer (VWR; Radnor, PA, USA) cells were filtered through a 40 μM cell strainer and grown on petri plates in DMEM containing 10% FBS and 30% L929 conditioned media for 7 days with fresh L929 containing media media being added on day 4 post plating. IL-4 (PeproTech; Rocky Hill, NJ, USA) was used at a concentration of 20ng/mL for *in vitro* experiments unless otherwise stated. Human U937 monocytic cells were a gift from Dr. Kristin Patrick and maintained in Roswell Park Memorial Institute 1640 Medium (RPMI 1640; Gibco, ThermoFisher Scientific, Waltham, MA, USA) supplemented with 10% FBS. To differentiate U937 monocytes into macrophages, cells were treated with 100ng/mL phorbol 12-myristate 13-acetate (PMA; AmBeed, Arlington Hts, IL, USA) for 48 hours. After that time period the media was changed, and cells were treated similarly as BMBMs. All cells were cultured at 37 °C with 95% humidity and 5% CO_2_.

### Lentiviral production

Twenty-four micrograms of each Lenti-CrispR-V2 plasmids containing a guide RNA against NIK and 72 μg of polyethyleneimine were used to transfect HEK293T cells. After 3 days of transfection, viral supernatant was harvested and filtered through a 0.45-μM syringe filter. After filtration, viral particles were concentrated 20-fold to 400 μl using Lenti-X Concentrator (Clontech, #631231, Mountain View, CA), and 100 μl of concentrated virus was used to infect cells. 1 μg/mL Puromycin (InvivoGen; San Diego, CA, USA) was used for selection until no non-transduced cells were alive.

### CRISPR-Cas9 gene knockout

U937 cells were transduced with a mixture of Lenti-CrispR-v2 virus’ carrying three gRNAs for each target. The gRNA sequences for human NIK were previously described (20, 21). Loss of NIK was confirmed by immunoblot of puromycin-resistant cells. Single colony cells were isolated by serial dilution. All experiments were repeated with at least three knockdown clones

### Seahorse Extracellular Flux Analysis

The Seahorse XFe96 Analyzer (Agilent; Santa Clara, CA, USA) was used to measure mitochondrial respiration. 5 × 10^4^ cells were plated per well in at least triplicate in 100 μL of DMEM media containing 10%FBS in an Agilent Seahorse XF96 Cell Culture Microplate. After allowing the cells to settle at room temperature for one hour, cells were moved to a 37°C incubator for 3 hours until stimulated overnight with 20 ng/mL IL-4. After 16 hours the cells were washed and replaced with XF assay medium (Base Medium Minimal DMEM supplemented with 200mM L-glutamine pH 7.4) and incubated with a 37°C incubator without CO2 for an hour before analysis. The Oxygen Consumption Rate (OCR) was measured after sequential addition of 25 mM glucose, 1.5 μM Oligomycin A (Sigma Aldrich; Saint Louis, MO, USA), 1.5 μM FCCP (Sigma Aldrich; Saint Louis, MO, USA) plus 1 mM Sodium Pyruvate (Gibco, ThermoFisher Scientific; Waltham, MA, USA), and 2.5 μM Antimycin A (Sigma Aldrich; Saint Louis, MO, USA) plus 1.25 μM Rotenone (Enzo Life Sciences, Farmingdale, NY, USA) following a previously published protocol (29).

#### NIK Inhibition

Same workflow as above with the exception of after allowing the cells to settle, DMSO or 5μM B022 (MedChem Express, Monmouth Junction, NJ, USA) were added to the cells and allowed to incubate for up to 5 hours before treating the cells overnight with 20ng/mL IL-4.

### RNA Sequencing

1 × 10^6^ NIK^WT^ and NIK^KO^ BMDMs were stimulated with 20ng/mL IL-4 or PBS for 6 hours before being lifted with PBS containing 1mM EDTA (Gibco, ThermoFisher Scientific; Waltham, MA, USA). Cells were spun down and washed once with PBS before being flash frozen in liquid nitrogen. Cell pellets were shipped on dry ice to Genewiz (South Plainfield, NJ, USA) where they isolated the RNA, constructed the sequencing library, and conducted the sequencing. RNA-sequencing data was analyzed using the Salmon pipeline (version 1.4.0-gompi-2020b) on the Texas A&M High Performance Research Computing Cluster. Salmon was used to align the results to the reference genome *Mus muculus* GRCm39. The data was then processed through DESeq2 (version 1.32.0) on R Studio where the annotated gene expression files were used to determine statistically significant changes in gene expression in stimulated NIK^KO^ BMDMs relative to stimulated NIK^WT^ BMDMs. Ingenuity Pathway Analysis software (IPA; QIAGEN, Germantown, MD, USA) was used to identify gene families and potential upstream regulators within the datasets. GraphPad Prism (San Diego, CA, USA) was used to make heatmaps. Raw and analyzed RNASeq datasets can be accessed at the NCBI GEO, accession number XXXXXX. (the GEO submission is pending and will be updated as soon as approved)

### Gene Expression

5×10^5^ NIK^WT^ and NIK^KO^ BMDMs were stimulated with 20ng/mL IL-4 for 6 hours before being lifted with PBS containing 1mM EDTA (Gibco, ThermoFisher Scientific; Waltham, MA, USA). Cells were spun down and washed once with PBS before total RNA was isolated from cells by Purelink™ RNA Mini Kit (Life Technologies). cDNA was synthesized from 1 μg total RNA using iScript reverse transcription supermix (Bio-Rad, Hercules, CA) following the manufacturer’s protocol. Quantitative RT-PCR was performed using Universal SYBR Green Supermix with StepOnePlus Real-Time PCR System (Applied Biosystems, Foster City, CA). Data was graphed as a 2^ΔΔ CT^ fold change with the gene expression data being normalized to the house keeping gene RPS18.

### Immunofluorescence

BMDMs were grown on 12mm coverslips at 6 × 10^5^ cells/well. Cells were treated with 200ng/ml LPS for 24 hours. After washing in PBS, cells were fixed with 4% paraformaldehyde for 20 min, permeabilized with 0.1% Triton X-100 in PBS for 5 min, blocked with PBS containing 5% FBS for 30 min, stained with primary antibodies for 1 hour, and stained with secondary antibodies for 1 hour. Cells were washed with PBS containing 5% FBS between each step Coverslips were mounted with Prolong Gold anti-fade reagent containing DAPI (Molecular Probes). Cells were imaged on an Olympus FV3000 confocal laser scanning microscope using a 100X oil-immersed objective.

### Mitochondrial isolation

2 × 10^6^ NIK^WT^ and NIK^KO^ BMDMs were plated in a 6 well plate and treated as described previously (22). At the time of harvest, cells were washed with PBS and scraped into a 1.5mL tube before spinning down. PBS was replaced with 0.5mL isolation buffer (320mM Sucrose, 1mM EDTA, 10mM HEPES, pH 7.2 with KOH and stored at 4°C) containing protease inhibitors (ThermoFisher Scientific; Waltham, MA, USA). While on ice, cells in isolation buffer were passed through a 27-gauge needle 10 times before spinning at 700xg for 8 minutes at 4°C. Supernatant was removed and placed into a new tube and kept on ice while the pellet was resuspended in 0.5mL isolation buffer with protease inhibitors before spinning at 700xg for 5 minutes at 4°C. Supernatant was removed and added with the previous supernatant, pellet contains nucleus, and the 1mL of supernatant was spun down at 17,000xg for 11 minutes at 4°C. After spinning the supernatant was placed into a new tube labeled cytoplasm and the pellet was resuspended in isolation buffer and spun down at 17,000xg for 11 minutes at 4°C. Following this last spin, the supernatant was discarded, and the mitochondrial pellet was resuspended in 40uL isolation buffer and protein concentration was measured by BCA assay using the Victor X3 microplate reader.

### Electron transport chain activity assays

Electron transport chain activity assays were obtained from Cayman Chemical Company (Ann Arbor, MI, USA) and manufacturer’s directions were followed with the following changes. 50uL of tube A was added to each well in the 96 well plate but activity buffer was used in place of the supplied mitochondria. We then added 5ug of mitochondrial protein (isolation described above) to a total volume of 20uL activity buffer per well. At this point each well should have 70uL per well before adding 30uL of tube B to start the reaction. Reactions were conducted at 25°C using an Epoch microplate reader with readings taken every 45 seconds for 20 minutes.

### ELISA

2 × 10^6^ NIK^WT^ or NIK^KO^ BMDMs were plated in a 6 well plate and allowed to adhere overnight. The following day the cells were treated with PBS or 20ng/mL IL-4 for 24 hours. Media was then removed from the cells, spun down, and froze at-20°C until ready for analysis (cells were used for other experiments). IL-10 ELISA matched antibodies were obtained from Tonbo Biosciences (San Diego, CA, USA) and the ELISA was carried out according to the manufacturer’s instructions. TMB Substrate was used to detect the peroxidase-labeled secondary antibody in the ELSIA kit (KPL, Gaithersburg, MD, USA). 100uL of undiluted media was used and ran in duplicate for each biological replicate.

### Macrophage Heptotaxis Assay

This assay was described previously (30). In brief, NIK^WT^ and NIK^KO^ BMDMs were stimulated with IL-4 (20ng/mL) for 24 hours before lifted with PBS containing 1mM EDTA (Gibco, ThermoFisher Scientific; Waltham, MA, USA). 75,000 DiO (ThermoFisher Scientific; Waltham, MA, USA) labeled BMDMs were plated on the insert of an 8.0μm pore, polyester membrane Transwell Permeable Support (Corning Incorporated; Corning, NY, USA) which was precoated for 3 hours at 37°C with 4ug/mL fibrinogen (Sigma-Aldrich; Saint Louis, MO, USA). After letting the cells settle for 3 hours, the media was removed and a Matrigel Matrix (50% Matrigel, 49% Culture media, 1% FBS) was added on top of the cell layer (Corning Incorporated; Corning, NY, USA). Once the Matrigel matrix solidified, a solution of 100nM

N-Formyl-Met-Leu-Phe [FMLP] (Sigma-Aldrich; Saint Louis, MO, USA) was added to the top of the Matrigel to induce a chemokine gradient. Cells were allowed to migrate for 24 hours before z-stack images were acquired on a Nikon A1R Confocal microscope (Nikon Instruments; Melville, NY, USA). Images were then processed in Imaris 9.5.0 (Oxford Instruments; Abingdon, Oxfordshire, UK) before counting the number of cells invaded. Data was graphed as a fold change to NIK^WT^ of the number of total cells that invaded.

### Macrophage Adhesion Assay

This assay was described previously (30). In brief, NIK^WT^ and NIK^KO^ BMDMs were stimulated with IL-4 (20ng/mL) for 24 hours before being lifted with PBS containing 1mM EDTA (Gibco, ThermoFisher Scientific; Waltham, MA, USA). Cells were diluted to a concentration of 1 million cells/mL and 50uL of cell solution was plated on a precoated 4ug/mL fibrinogen and 0.5% polyvinyl alcohol (Sigma-Aldrich; Saint Louis, MO, USA) 96-well plate. Cells were plated with technical replicates and allowed to sit for 30 minutes in a 37°C tissue culture incubator. Cells were then washed once with PBS before staining the adhered cells with the CyQUANT DNA stain according to the manufacturer’s instructions (Invitrogen, ThermoFisher Scientific; Waltham, MA, USA). Fluorescence signal was measured on the Victor X3 microplate reader and cell number was determined through a standard curve of cells.

### Macrophage Phagocytosis Assay

NIK^WT^ and NIK^KO^ BMDMs were plated subconfluently in a 6-well plate before being stimulated with IL-4 (20ng/mL) for 24 hours. 10^8^ 1um carboxylate microspheres (Polysciences Inc; Warrington, PA, USA) were coated in a 50% mouse serum (MP Biomedicals; Irvine, CA, USA) solution and incubated for 30 minutes at 37°C. The solution was then diluted to 5% serum before being added to cells and allowed to incubate for 2 hours. After, cells were washed twice with PBS before fixing with 4%PFA. Once the PFA was washed, cells were stained with Phalloidin and Hoechst (Invitrogen, ThermoFisher Scientific; Waltham, MA, USA) and imaged on a Nikon Eclipse Ti microscope.

### Statistical Analysis

Statistical analyses were carried out using GraphPad Prism (San Diego, CA, USA), and details can be found in the figure legends where applicable. The data presented herein were considered statistically significant at p◻< ◻0.05.

## Results

### NIK regulates macrophage metabolism by shaping mitochondrial respiratory capacity and OXPHOS complex activity

To investigate how NIK regulates innate immune cell function, we examined metabolic shifts during the polarization of bone marrow-derived macrophages (BMDMs) obtained from wild-type (NIK^WT^) and NIK knockout (NIK^KO^) mice. After differentiation in L929-conditioned media for 7 days, we noted that M0 macrophages from NIK^KO^ mice displayed reduced basal and maximal respiration, as well as impaired mitochondrial spare respiratory capacity (SRC) (**Figure 1A**). Upon treatment with interleukin-4 (IL-4) to stimulate an M2-like phenotype (M(IL-4)), NIK^KO^ BMDMs were unable to increase basal and maximal respiration, abrogating SRC and indicating a pronounced and persistent defect in oxidative metabolism (**Figure 1B**). Decreased maximal respiration and SRC were also observed in M0 and M(IL-4) NIK CRISPR knockout (NIK^-/-^) human monocytic U937 cells after their differentiation into macrophages (**Figure 1C-E**). Similar decreases in basal and maximal respiration were observed in NIK conditional knockout BMDMs (LysMCre; NIK^fl/fl^) and in NIK^WT^ BMDMs treated with the NIK inhibitor B022 (31) (**Supplemental Figure 1)**. Taken together, these data indicated that NIK regulates oxidative metabolism by controlling OXPHOS and maximal respiration.

**Figure 1:**
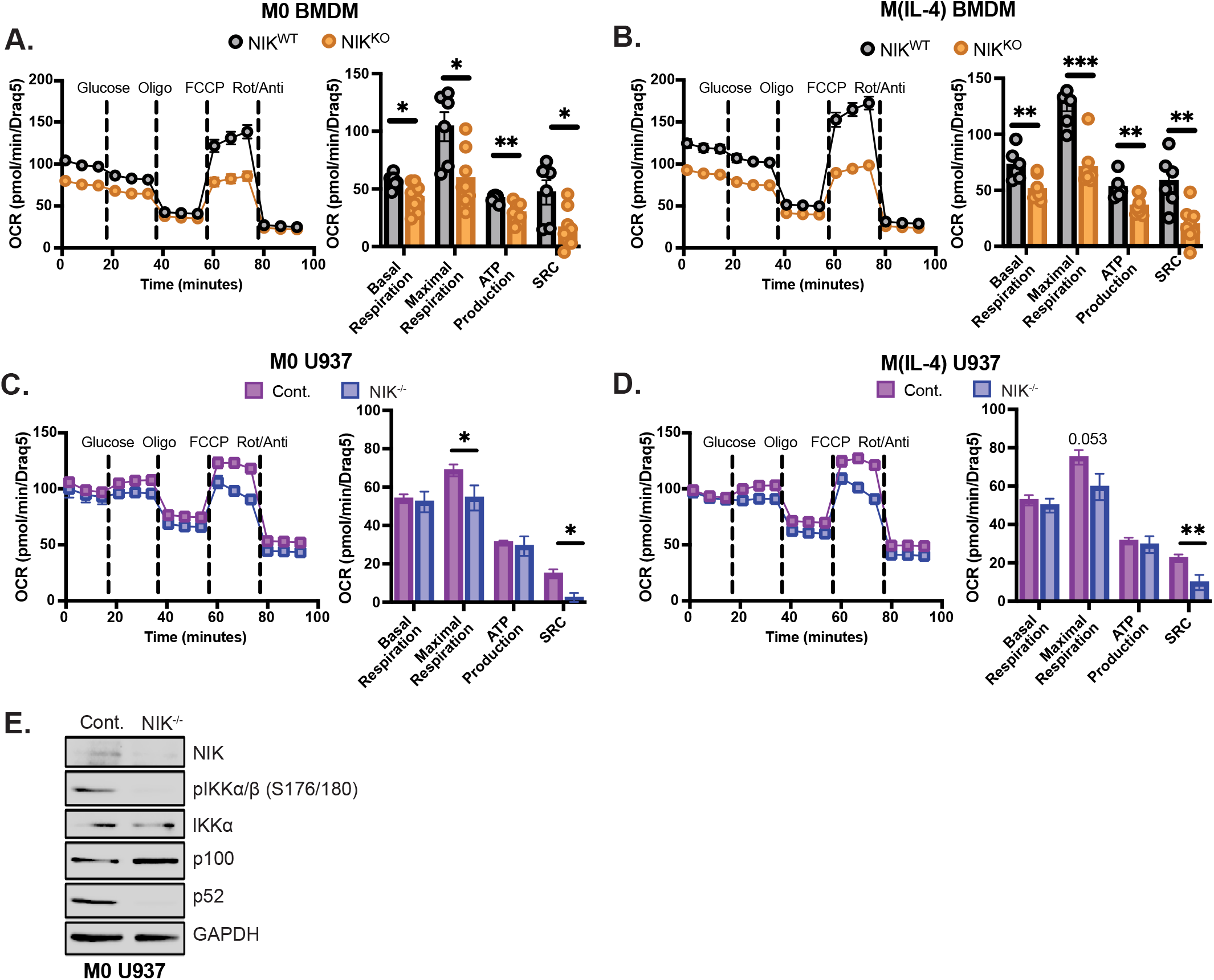
NIK regulates oxidative metabolism in macrophages. Seahorse extracellular flux analysis investigating changes in the oxygen consumption rate (OCR) of basal (M0) NIK^WT^ and NIK^KO^ BMDMs **(A)** or after an overnight treatment with IL-4 M(IL-4) **(B)**. Line graphs are combined technical (n > 23) and biological replicates with bar graphs representing the averaged technical replicate value for each animal (n > 6 biological replicates). Data are mean +/-SEM and statistics are a Student’s T-test *p < 0.05, **p < 0.01, ***p < 0.001. Seahorse extracellular flux analysis showing **(C)** OCR readouts of basal (M0) and **(D)** IL-4 treated (M(IL-4)) control and NIK^-/-^ U937 cells. Graphs represent 3 independent experiments with treatment conditions ran in at least technical triplicates (n = 3 single cell isolated clones). Data are mean +/-SEM and statistics are a Student’s T-test. *p < 0.05, **p < 0.01. **(E)** Representative western blot verifying loss of NIK in U937 cells using the empty CRISPR vector, LCV2, as a control. Samples were treated with 200ng/mL LPS and 10μM MG132 for 4 hours before extracting protein with RIPA buffer.

To gain insight into the underlying OXPHOS deficit in NIK^KO^ BMDMs, we performed RNA sequencing on M0 and M(IL-4) skewed BMDMs. We did not observe significant changes in the expression of genes involved in glycolysis, TCA cycle, pentose phosphate pathway, glucose transporters, or electron transport chain subunits (**Supplemental Figure 2**), indicating that loss of respiratory capacity in NIK^KO^ BMDMs occurs at a post-transcriptional level. However, we did note a significant increase in mitochondrial superoxide production in both M0 and M(IL-4) NIK^KO^ BMDMs compared to NIK^WT^ (**Figure 2A**,**B**), indicating enhanced electron leak from inefficient respiratory complexes. Mitochondrial reactive oxygen species (mROS) can be produced from OXPHOS complexes I, II, and III, so we next performed enzymatic assays with isolated mitochondria to assess respiratory complex efficiency. Consistent with our mROS data, we observed that NIK^KO^ BMDMs exhibited impaired activity of Complexes I and II compared to NIK^WT^ BMDMs (**Figure 2C,D**). This apparent electron transport chain defect correlated with defects in other markers of mitochondrial fitness, including increased numbers of mitochondria in the NIK^KO^ BMDMs with reduced membrane potential (**Supplemental Figure 3A-C**) and a trend toward increased mitochondrial size (**Supplemental Figure 3E-G**). Complexes III-V exhibited no changes in activity between NIK^WT^ and NIK^KO^, regardless of treatment with IL-4 (**Figure 2E-G**). Similar to our previous findings in cancer cells (21, 22), NIK localized to the mitochondria after treatment with IL-4 (**Supplemental Figure 3D**). These data suggest that mitochondria-associated NIK controls oxidative metabolism and electron transport chain function through OXPHOS Complexes I and II.

**Figure 2:**
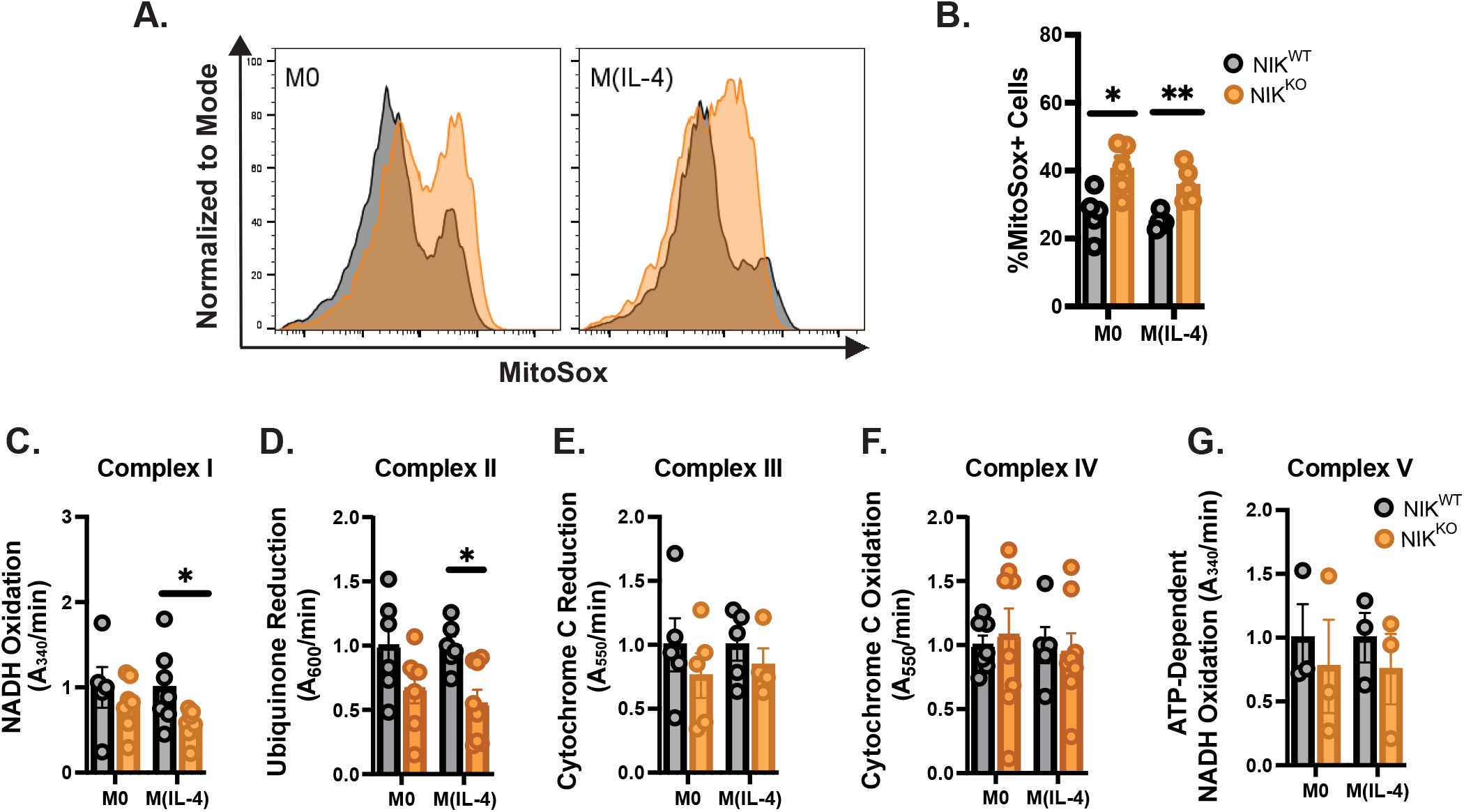
NIK deficient macrophages display elevated mROS and impaired OXPHOS Complex I and II activity. (**A,B)** Representative histograms and quantification of Mitosox levels in NIK^WT^ and NIK^KO^ BMDMs post 6-hour treatment with IL-4 (n = 5 biological replicates). Statistics are a Student’s T-test. *p < 0.05, **p < 0.01. Electron transport chain activity assays for Complex I **(C)**, Complex II **(D)**, Complex III **(E)**, Complex IV **(F)**, and Complex V **(G)** using isolated mitochondria from basal or 24 hour IL-4 treated NIK^WT^ and NIK^KO^ BMDMs. Graphs represent biological replicates (n > 3 biological replicates ran in at least duplicate). Statistics are a Student’s T-test. *p < 0.05.

**Figure 3:**
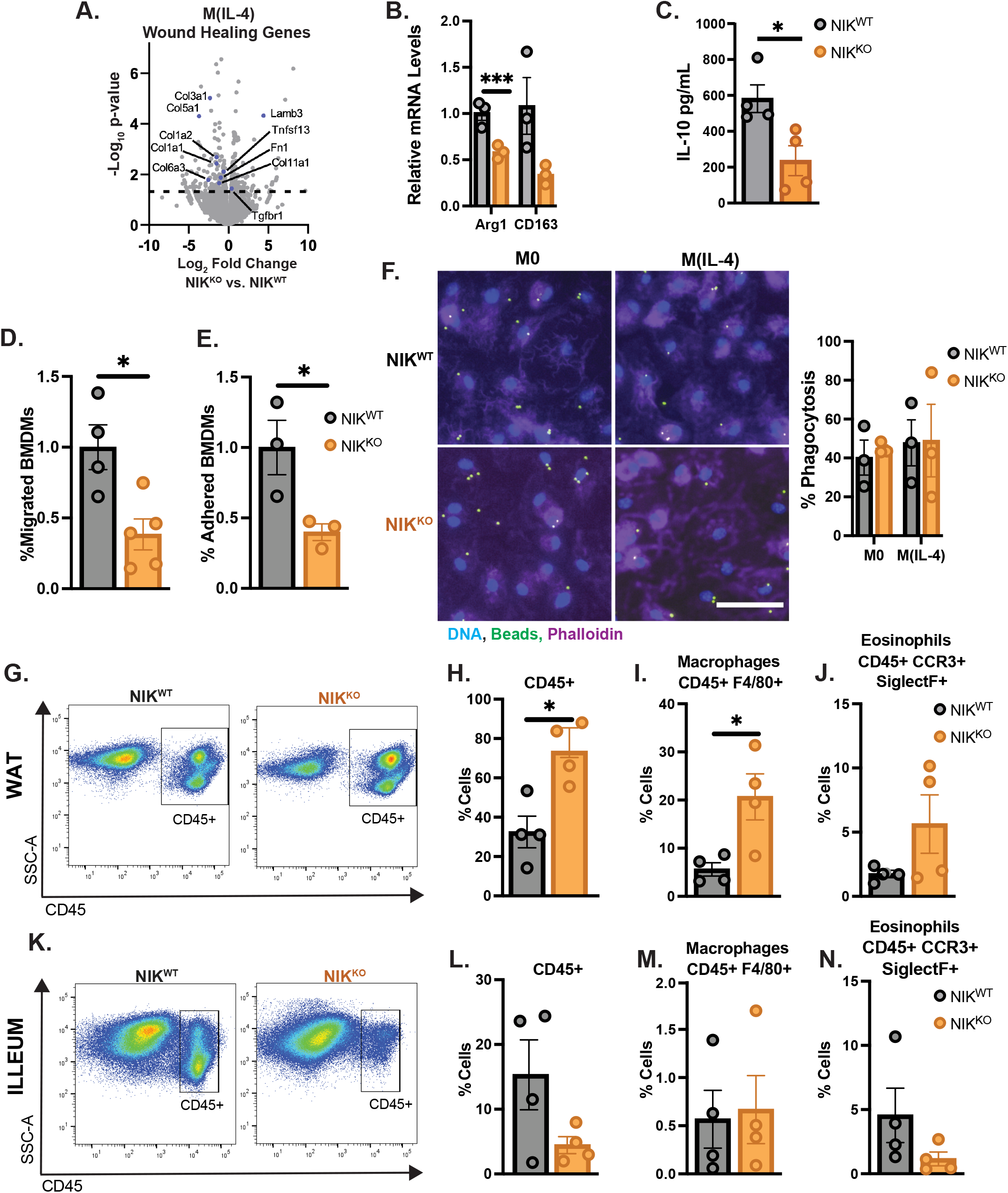
NIK^KO^ macrophages fail to polarize to an M(IL-4) phenotype *in vitro* and *in vivo*. **(A)** RNA-sequencing data of classical pro-repair pathways in NIK^KO^ macrophages after 6 hours of IL-4 treatment. Volcano plots represent 2 biological replicates with a Log2Fold change of NIK^KO^ BMDMs compared back to NIK^WT^ of the same treatment. Statistics were calculated by DESeq2 and genes above the dotted line are deemed significant for having a-log(p-value) of greater than 1.3. **(B-C)** The trends in RNA-sequencing were confirmed by assessing IL-4 response genes in by RT-qPCR **(B)** and validating IL-10 levels by ELISA post 24hr IL-4 treatment in NIK^WT^ and NIK^KO^ macrophages **(C)** (n > 3 biological replicates ran in triplicates). Statistics are a Student’s T-test *p < 0.05, ***p < 0.001. NIK^WT^ and NIK^KO^ BMDMs were stimulated for 24 hours with IL-4 before undergoing the following assays: **(D)** Cell invasion assay was performed with media containing the chemoattractant FMLP. After 24 hours invaded cells were calculated percent migrated compared to NIK^WT^ (n > 4 biological replicates). Statistics are a Student’s T-test *p < 0.05. **(E)** Cells were plated on a PVA coated 96 well plate and allowed to settle for 30 minutes before washing with PBS and quantifying percent attached cells via CyQUANT DNA staining and comparing the NIK^KO^ value back to NIK^WT^ (n = 3 biological replicates). Statistics are a Student’s T-test *p < 0.05. **(F)** Representative fluorescent images of phagocytosis assay. Graph represents % BMDMs containing beads (n = 3 biological replicates). Statistics are a Student’s T-test. *In vivo* analysis of NIK^WT^ and NIK^KO^ mice investigating immune cells in the WAT **(G-J)** and ileum **(K-N)** gated through single and live cells. Quantification of biological replicates analyzing total immune cells, macrophages, and eosinophils in WAT (top) and ileum (bottom) is graphed as a percent of CD45+ parent gate (n = 4). Statistics are a Student’s T-test. *p < 0.05.

### NIK^KO^ Macrophages Fail to Polarize to an M(IL-4) Phenotype *in vitro* and *in vivo*

To investigate the downstream pathways impacted by the dysregulated metabolism observed in the absence of NIK, we performed transcriptome analysis of NIK^WT^ and NIK^KO^ BMDMs, and evaluated transcriptional changes in macrophage phenotype signatures. We observed that there were significant defects in M2-related wound healing gene signatures in the NIK^KO^ BMDMs compared to NIK^WT^ in response to IL-4 treatment (**Figure 3A**). The RNA-sequencing data were verified via qPCR for changes in expression of typical IL-4 response genes, such as Arginase 1 (Arg1) and CD163, and a significant decrease in expression was observed in Arg1 in the NIK^KO^ BMDMs compared to NIK^WT^ (**Figure 3B**). Consistent with this finding, we observed a significant decrease in IL-10 secretion to the cell culture media in response to IL-4 treatment (**Figure 3C**). Additionally, after IL-4 treatment, we observed that NIK^KO^ BMDMs harbored functional defects in migration and adhesion (**Figure 3D,E**), although their phagocytic capabilities remained intact (**Figure 3F**). Taken together with the metabolic findings discussed above, these data suggest that NIK^KO^ BMDMs fail to fully polarize into an M(IL-4) phenotype due to a deficiency in OXPHOS activity.

Eosinophils have been shown to secrete IL-4 and promote an M(IL-4) macrophage phenotype *in vivo* (32–35). Increased peripheral and tissue eosinophil numbers have been previously described in NIK^KO^ mice and are associated with dermatitis and esophageal eosinophilia (25, 26). Because little is known about the role of NIK in macrophages, we investigated the relationship between the accumulated eosinophils and their effect on macrophage polarization in visceral white adipose tissue (WAT) deposits, which are known to have low eosinophil numbers and high levels of macrophages, along with the small intestine, which has high eosinophil numbers and low levels of macrophages (33, 35, 36). It was previously reported that NIK^KO^ mice exhibit decreased body weight compared to NIK^WT^ controls, which also correlates with decreased fat mass and smaller adipocytes (37). We found that NIK^KO^ mice had a significant accumulation of lineage positive immune cells within the WAT compared to NIK^WT^ littermates and further analysis of the immune populations revealed a significant increase in macrophages and a similar trend in eosinophils (**Figure 3G-J**). In contrast, total immune cells, macrophages, and eosinophils were either unchanged or decreased in the NIK^KO^ ileum compared to NIK^WT^ (**Figure 3K-N**). Taken together, the changes in immune populations within WAT suggest persistent inflammation due to high levels of macrophages and eosinophils.

### NIK^KO^ Mice Have Increased Levels of Pro-Inflammatory Myeloid Cells in both the Blood and Bone Marrow

Given the changes in immune populations observed in the WAT and small intestine in NIK^KO^ mice, we performed high dimensional multiparameter flow cytometry and standard gating to phenotype circulating immune cell populations. Consistent with prior findings and roles for NIK in B cell development and function (15, 16, 24, 38), we observed markedly reduced CD19+ B lymphocytes, but not CD3+ T cells, in the blood of NIK^KO^ mice (**Figure 4A-B**). We also observed a large increase in the percentage of CD11b+ myeloid cells, which was characterized by a shift from neutrophils to eosinophils and Ly6C^hi^ inflammatory monocytes (**Figure 4C-F**). Next, using t-Distributed Stochastic Neighbor Embedding (tSNE) analysis to investigate myeloid immune marker expression more broadly, we confirmed that NIK^KO^ mice contain dramatically more inflammatory eosinophils (**Figure H, green dashed line**) (25, 26). These myeloid cells were characterized as side scatter high (SSC-A), CD11b+, Ly6C+, Ly6G^lo^, F4-80+, and high expression of TNFα at rest (**Figure 4G-I**). We also observed that NIK^KO^ mice possess elevated TNFα+, Ly6C^hi^ inflammatory monocytes in the blood at rest (**Figure 4G-I, black dashed line**). Ex vivo LPS stimulation of blood leukocytes revealed a greater enhancement of CD11b+ Ly6C^hi^ inflammatory monocytes in NIK^KO^ mice compared to NIK^WT^ (**Figure 4J**). Four hours after LPS stimulation, CD11b+ Ly6C^hi^ monocytes from NIK^KO^ mice produced very high levels of intracellular TNFα relative to NIK^WT^ monocytes **(Figure 4J,K,L**). Overall, these results reveal that NIK^KO^ mice have a significant increase in blood myeloid cells that are both basally producing pro-inflammatory cytokines and hyper-responsive to innate immune stimuli, suggesting unresolved systemic inflammation.

**Figure 4:**
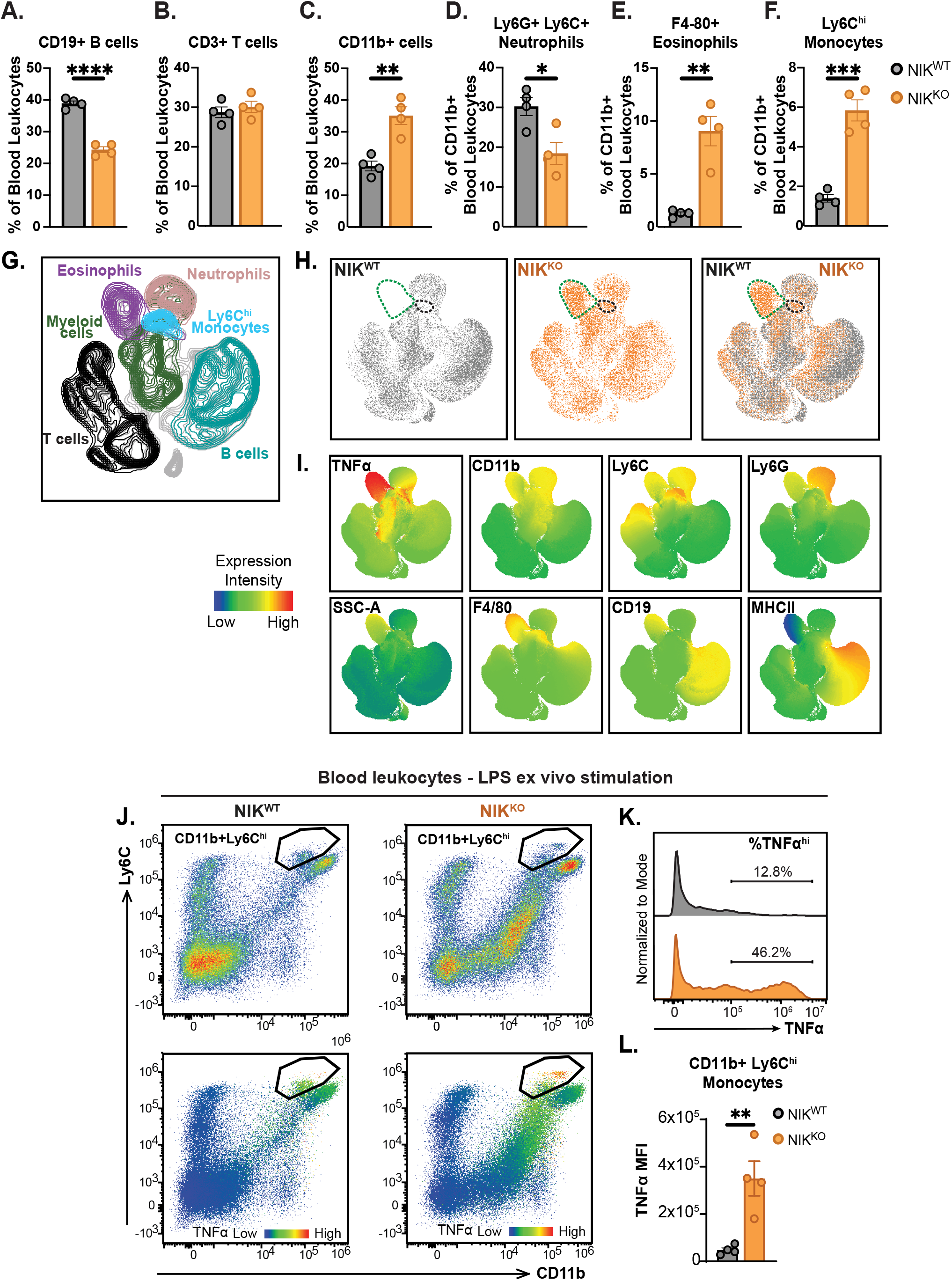
NIK^KO^ mice exhibit increased pro-inflammatory myeloid cells in the blood at baseline and after innate immune stimulation. Blood was harvested from NIK^WT^ and NIK^KO^ mice and subjected to red blood cell lysis with ACK buffer. Total blood leucocytes were quantified by flow cytometry using markers of B cells **(A)**, T cells **(B)**, CD11b+ myeloid cells **(C)**, neutrophils **(D)**, eosinophils **(E)**, and monocytes **(F)**. (n = 4 biological replicates). Statistics are a Student’s T-test. *p < 0.05, **p < 0.01, ***p < 0.001, ****p < 0.0001. **(G-I)** T-SNE clustering of NIK^WT^ and NIK^KO^ blood leukocytes representing 4 biological replicates. **(J-L)**. Blood leukocytes were stimulated with LPS ex vivo and TNFα mean fluorescence intensity (MFI) levels were quantified. Statistics are a Student’s T-test. **p < 0.01.

Given the role that NIK plays a role in hemopoietic stem cell differentiation (17, 18), we next investigated the bone marrow of NIK^KO^ mice for defects in immune populations. We found that NIK^KO^ mice have a significant increase in the total number of bone marrow cells (**Figure 5A**), which are primarily derived from increased CD45+ lineage positive cells (**Figure 5B,C**). Analysis of CD45+ cells in NIK^KO^ mice showed a significant increase in granulocytes compared to blast cells, monocytes, and lymphocytes (**Figure 5D-G**). Moreover, we found that the increase in granulocytes is due to an increase in eosinophils as opposed to neutrophils (**Figure 5H,I**). Consistent with our blood findings, we observed that monocytic cells were slightly elevated (**Figure 5J**), total T cells were unchanged (**Figure 5K**), and B cells were significantly decreased (**Figure 5L**). These data suggest that dysregulated terminal differentiation of cell populations in circulation and the bone marrow stem from an enrichment of low proliferative cells within the stem cell niche of the bone marrow in NIK^KO^ mice (**Supplemental Figure 4**). Taken together, our findings demonstrate that NIK^KO^ mice exhibit a systemic skewing toward myeloid and pro-inflammatory phenotypes in the bone marrow, circulation, and peripheral tissues, driven at least in part by dysregulated mitochondrial homeostasis and oxidative metabolism (**Figure 5M,N**).

**Figure 5:**
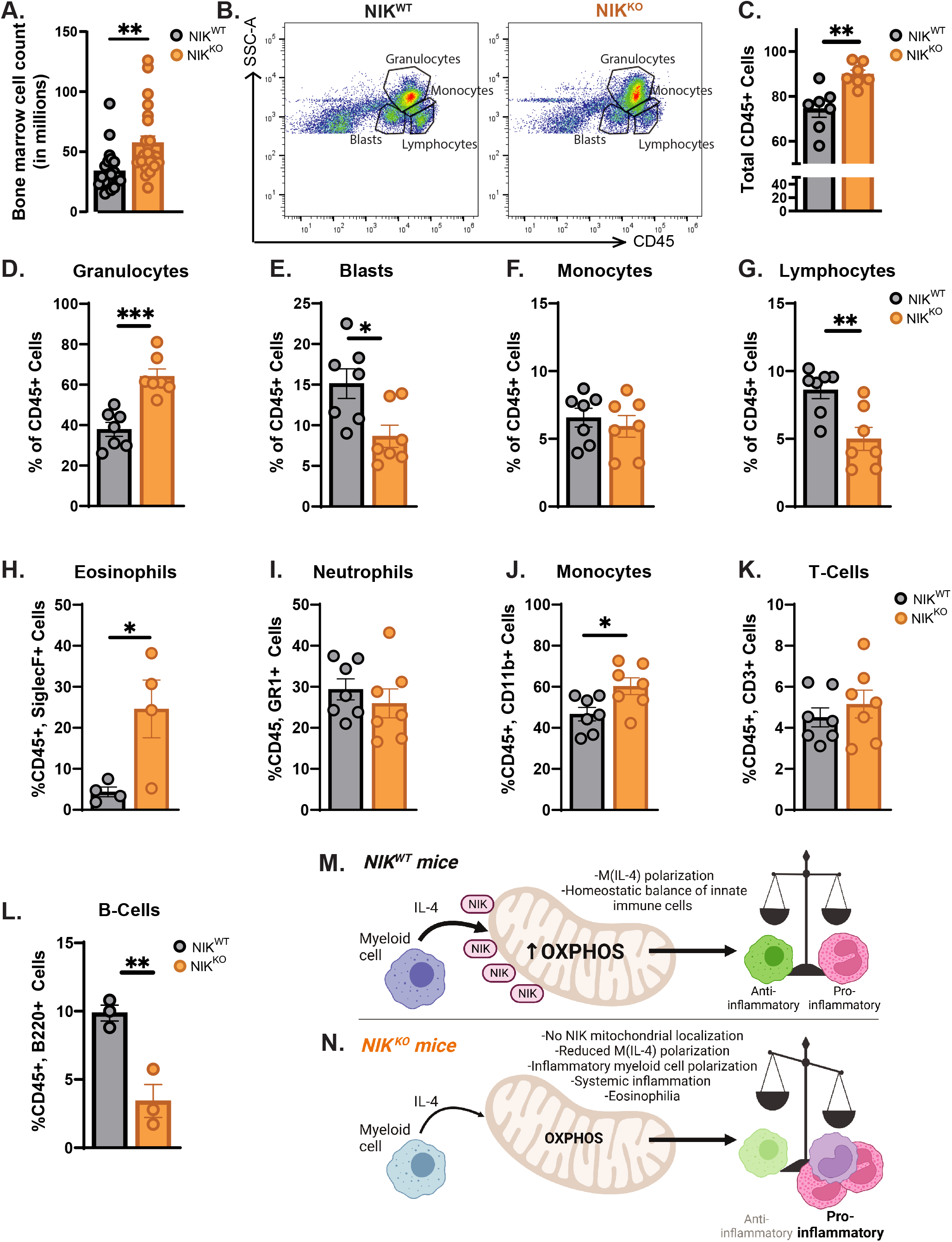
NIK^KO^ mice exhibit myeloid skewing in the bone marrow. **(A)** Total bone marrow cell count after ACK red blood cell lysis (n = 23 biological replicates). Statistics are a Student’s T-test **p < 0.01. **(B**,**C)** Quantification and flow cytometry plot of lineage positive (CD45+) cells in NIK^WT^ and NIK^KO^ mouse bone marrow (n = 7). Statistics are a Student’s T-test **p < 0.01. **(D-G)** Quantification of different bone marrow cell populations in NIK^WT^ and NIK^KO^ bone marrow based off of Side Scatter Area (SSC-A) and CD45 (n = 7). **(H-L)** Quantification of specific immune populations in the CD45+ parent gate utilizing the indicated surface markers. Statistics are a Student’s T-test. *p < 0.05, **p < 0.01, ***p < 0.001. Model showing the role of NIK in WT myeloid cells promoting oxidative metabolism and having a homeostatic balance of innate immune cells **(M)** and the consequences of NIK loss resulting in decreased oxidative metabolism and systemic inflammation **(N)**. Model Created with BioRender.com.

## Discussion

Although prior studies have established a critical role for NIK in controlling adaptive immunity and development (15, 16, 23, 24, 38), much less is known about whether and how NIK regulates innate immune functions. While we have demonstrated that NIK acts as a stress sensor to regulate both basal and stress-induced mitochondrial metabolism to promote glioblastoma pathogenesis (22), it was not clear whether NIK played similar roles in non-cancerous immune cells. In this study, we identify a previously unreported requirement for NIK in regulating immunometabolic switches in myeloid cells to promote their acquisition of a pro-repair, anti-inflammatory immune phenotype. Specifically, we observed that BMDMs isolated from NIK^KO^ mice exhibited impaired OXPHOS, with a significant reduction in the maximal respiratory and spare respiratory capacities, and the inability to fully acquire an M(IL-4) polarization state, as seen by a failure to induce M2 markers (see **Figure 3**). Similarly, decreased mitochondrial respiration was also observed in BMDMs isolated from mice with a conditional deletion of NIK (LysMCre NIK^fl/fl^, **see Supplemental Figure 1**), as well as NIK knockout human monocytic U937 cells differentiated into macrophages, indicating cell intrinsic mitochondrial functions for NIK. Furthermore, we observed that pharmacological inhibition of NIK in BMDMs also impaired mitochondrial respiration, indicating a requirement for the NIK kinase functionality in innate immune cell metabolic regulation. On a systemic level, we observed that NIK^KO^ mice exhibit a skewing toward pro-inflammatory myeloid phenotypes in the bone marrow, blood, and adipose tissue, characterized by increased numbers of macrophages, eosinophils, and monocytes expressing high levels of TNFα. Since both cell-intrinsic and cell-extrinsic NIK deficiency affects systemic inflammation (25), future studies are required to address whether the phenotypic changes observed in myeloid cell populations in NIK^KO^ mice are controlled cell-autonomously.

Our findings are the first to demonstrate a requirement for NIK in myeloid cell mitochondrial metabolism and macrophage polarization through regulation of electron transport chain function. Moreover, while we noted a decrease in bone marrow and blood CD45+ B lineage cells in NIK^KO^ mice as previously reported (38), we demonstrate a significant increase in eosinophils and CD11b+, Ly6C^hi^ monocytes expressing elevated TNFα at baseline and after LPS challenge. These findings suggest that previously reported hyper-eosinophilia and dermatitis in NIK^KO^ mice are associated with an inability of eosinophils to induce OXPHOS in macrophages to resolve inflammation (25, 26). Interestingly, development of eosinophilia in NIK^KO^ mice was shown to be independent of IKK/NF-κB transcriptional activity (25). This is consistent with our RNA-seq data confirming no significant transcriptional alterations in metabolic or macrophage polarization profiles in NIK^KO^ BMDMs, and suggests that the regulation of macrophage mitochondrial metabolism and TCA complex activity does not occur at a transcriptional level. Moreover, our findings are consistent with studies demonstrating that control of oxidative metabolism and mitochondrial dynamics in GBM cells (21, 22), glycolytic metabolism in T cells (23), and ATP production and respiratory capacity in B cells (24) are NIK dependent and independent of IKK/NF-κB dependent transcriptional regulation.

The most prominent reported phenotypes of NIK^KO^ mice, as well as mice harboring a spontaneous *NIK* mutation (NIK^aly/aly^), are their lack of lymph nodes and Peyer’s patches, aberrant splenic and thymic architecture, leading to profound defects in adaptive immunity and increased susceptibility to infection (15, 39, 40). Primary Immunodeficiency Disease (PID) patients who harbor loss-of-function mutations in *NIK* have remarkably similar clinical manifestations to NIK^KO^ mice, including lymphopenia, abnormal splenic architecture, and markedly reduced immunoglobulin levels leading to recurrent infections (15, 39–42). PID resulting from NIK mutation has been largely attributed to aberrant NF-κB transcriptional regulatory activity leading to adaptive immunodeficiency. How loss of NIK function alters innate immune cell development and function in patients with PID is unclear, and the impact of NIK dysregulation on immunometabolic homeostasis in PID has not been previously considered. We have recently observed that NIK^KO^ mice exhibit altered systemic metabolism characterized by an increase in aerobic glycolysis and impaired respiration (37), which is similar to patients with primary mitochondrial diseases (MD) caused by mutations in OXPHOS genes. Patients with MD often experience recurrent infections and display adaptive immunodeficiency (43, 44), paralleling those with NIK loss-of-function PID. We and others have found that mouse models of human mitochondrial disease display pronounced myeloid skewing in the bone marrow and blood, characterized by high levels of inflammatory neutrophils and monocytes (45, 46). Moreover, we observed that the Polymerase gamma (POLG) ‘mutator’ model of mitochondrial DNA disease exhibits OXPHOS deficits and a switch to glycolytic metabolism, which drive enhanced pro-inflammatory phenotypes, macrophage polarization, and tissue damage (46). Myeloid phenotypes in these mitochondrial mutant mice resemble findings here with NIK^KO^ mice. Taken together, these findings suggest that mitochondrial and metabolic dysfunction leading to innate immune rewiring could be a common feature of patients with NIK-deficient PID, Leigh Syndrome or other primary mitochondrial diseases. Future research to more thoroughly investigate these emerging links is warranted.

In conclusion, our findings elucidate a previously unrecognized role for NIK as a molecular rheostat that shapes the metabolic status, differentiation, and inflammatory phenotypes of myeloid cells in vitro and in vivo. Moreover, our work identifies NIK as a key factor governing innate immune cell mitochondrial function and suggests that metabolic dysfunction may be an important contributor to inflammatory and/or immunodeficiency diseases caused by aberrant NIK expression or activity. Our findings have important implications for human disease, as the bioenergetic pathways underlying immune cell phenotype and function are increasingly recognized as important nodes for therapeutic intervention in immune and inflammatory disorders. Moreover, our results provide a strong rationale for future analysis of overlapping phenotypes in inborn errors of immunity and inborn errors of metabolism and suggest that strategies aimed at boosting mitochondrial function may help rebalance the immune system in primary immunodeficiency caused by NIK loss of function.

## Acknowledgements

We thank Dr. Kristin Patrick for providing U937 cells.

## Disclosures

The authors declare no financial conflicts of interest.

---

**Supplemental Figure 1: Conditional knockout and chemical inhibition of NIK reduce maximal respiration**. Seahorse extracellular flux analysis investigating changes in the oxygen consumption rate (OCR) of basal (M0) LysMCre;NIK^+/+^ and LysMCre;NIK^fl/fl^ BMDMs **(A)** or after an overnight treatment with IL-4 M(IL-4) **(B)**. Line graphs are combined technical (n > 19) and biological replicates with bar graphs representing each animal (n = 5 biological replicates). Data are mean +/-SEM and statistics are a Student’s T-test *p < 0.05, **p < 0.01. **(C)** Representative western blot verifying loss of NIK in BMDMs from LysMCre mice. Samples were treated with 20ng/mL IL-4 and 10μM MG132 for 4 hours before extracting protein with RIPA buffer. Seahorse extracellular flux analysis showing **(D)** OCR readouts of IL-4 treated (M(IL-4)) NIK^WT^ BMDMs pretreated for 5 hours with 5μM B022 or vehicle (DMSO). Line graphs are combined technical (n > 20) and biological replicates with bar graphs representing each animal (n > 5 biological replicates). Data are mean +/-SEM and statistics are a Student’s T-test with pvalues shown on samples close to significant **p < 0.01. **(E)** Representative western blot showing inhibition of NIK post 5μM B022 treatment. Samples were pretreated with B022 for 5 hours before treatment with 20ng/mL IL-4 for 4 hours and extracting protein with RIPA buffer.

**Supplemental Figure 2: NIK**^**KO**^ **macrophages have no significant transcriptional changes in metabolic genes**. RNA-sequencing heatmaps represent 2 biological replicates with a Log2Fold change post 6 hour IL-4 treatment over baseline in NIK^WT^ **(A)** and NIK^KO^ BMDMs **(B)** probing for changes in genes involved in glycolysis, TCA cycle, pentose phosphate pathway, and glucose transporters. RNA-sequencing heatmaps of NIK^KO^ macrophages compared to NIK^WT^ at baseline **(C)** or after 6 hour treatment of IL-4 **(D)** representing changes in gene expression of electron transport chain complex subunits. Statistics were calculated by DESeq2 and -log10 (p value) of 1.3 represents a p value of 0.05.

**Supplemental Figure 3: Mitochondrial homeostasis is disrupted due to a loss of NIK. (A)** 24 hour IL-4 stimulated BMDMs were costained with Mitotracker green and Mitotracker red to assess mitochondrial membrane potential via flow cytometry as a marker of mitochondrial functionality. Cells with low mitochondrial membrane potential were termed “dysfunctional” **(B)** and those with high mitochondrial membrane potential “functional” (**C)**. Statistics are a Student’s T-test. *p < 0.05 (n = 5 biological replicates). **(D)** Representative western blots from human U937 cells showing NIK localization to a mitochondrial enriched fraction post 0-4 hours of IL-4 treatment along with mitochondrial fractionation controls. **(E)** Isolated mitochondria were analyzed by flow cytometry against standard sized beads to calculate mitochondrial size with the quantification of mitochondrial size based on the standard curve of the beads **(F)** (n = 4 biological replicates). Statistics are a Student’s T-test and pvalues are shown. **(G)** Representative immunofluorescent images characterizing the mitochondrial network in NIK^WT^ and NIK^KO^ macrophages at baseline using antibodies against HSP60 (green), mitochondrial DNA (red), TFAM (purple), and DAPI (blue).

**Supplemental Figure 4: Lineage negative bone marrow cells are enriched for low proliferative cells due to a loss of NIK**. Gating scheme for stem cell analysis in NIK^WT^ **(A)** and NIK^KO^ **(B)** bone marrow. Quantification of the stem cell compartment probing for differences in the LSK populations (early lymphoid progenitor) **(C)**, MPP (multi potent progenitor) **(D)**, LT-HSC (long term hemopoietic stem cell) **(E)**, and ST-HSC (short term hemopoietic stem cell) **(F)**. Cell populations (n ≥ 4 biological replicates). Statistics are a Student’s T-test. *p < 0.05, **p < 0.01.

## Notes

**Funding:** This work was supported by NIH grants R01HL148153 to A.P.W. and R01NS082554 to R.S. Additional support was provided by a pilot award from the Texas A&M School of Medicine and an X-Grant from the Texas A&M University Office of the Vice President for Research.

### Competing Interest Statement

The authors have declared no competing interest.

### Summary of Updates

This version corrects errors in the text, includes new data in multiple figures, and has an expanded discussion section.

