## Supplemental Figure 1 for "NF-κB-Inducing Kinase (NIK) Governs the Mitochondrial Respiratory Capacity, Differentiation, and Inflammatory Status of Innate Immune Cells"

### Figure S1

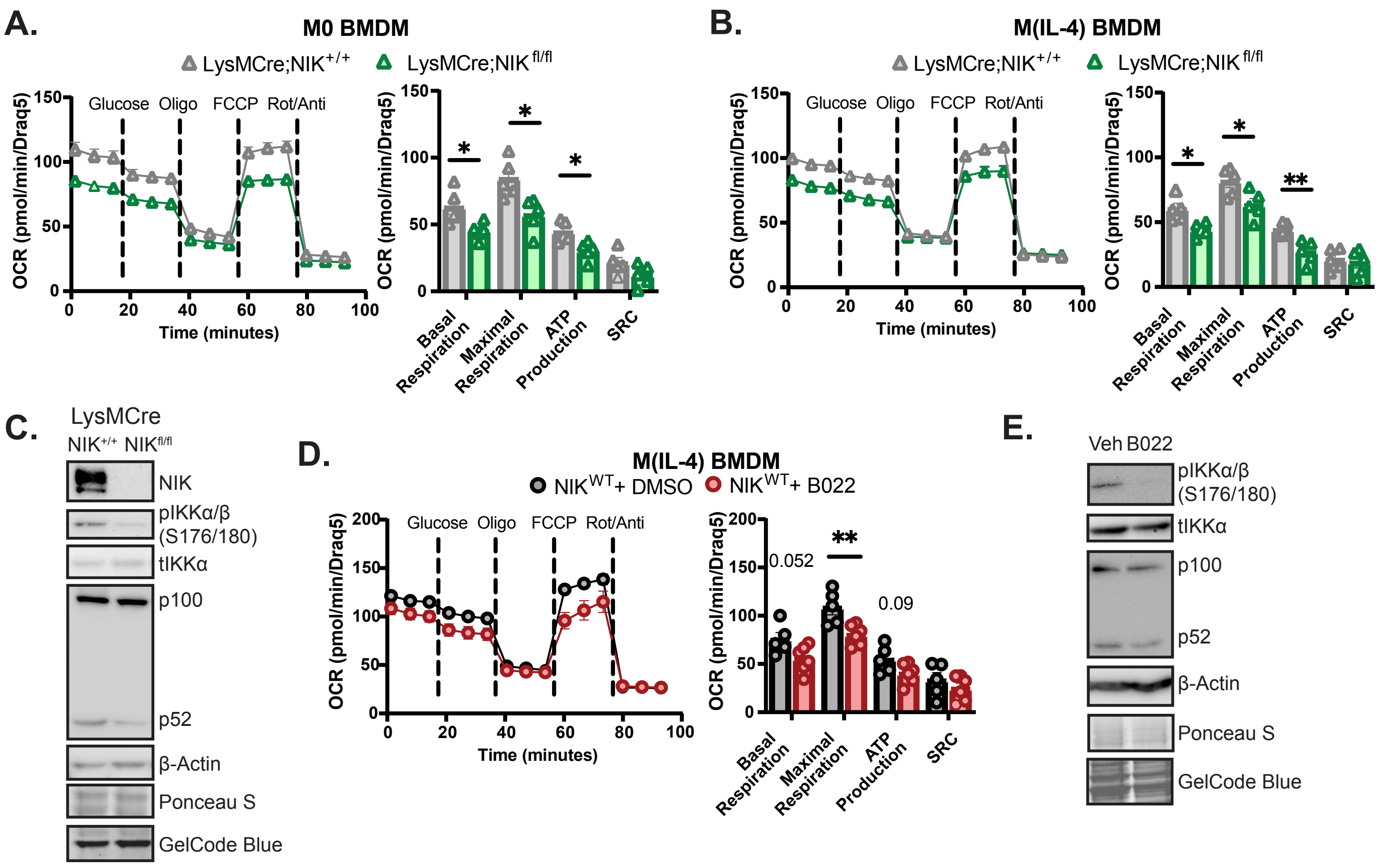


**Supplemental Figure 1: Conditional knockout and chemical inhibition of NIK reduce maximal respiration.** Seahorse extracellular flux analysis investigating changes in the oxygen consumption rate (OCR) of basal (M0) LysMCre;NIK+/+ and LysMCre;NIKfl/fl BMDMs (**A**) or after an overnight treatment with IL-4 M(IL-4) (**B**). Line graphs are combined technical (n > 19) and biological replicates with bar graphs representing each animal (n = 5 biological replicates). Data are mean +/- SEM and statistics are a Student's T-test *p < 0.05, **p < 0.01. (**C**) Representative western blot verifying loss of NIK in BMDMs from LysMCre mice. Samples were treated with 20ng/mL IL-4 and 10μM MG132 for 4 hours before extracting protein with RIPA buffer. Seahorse extracellular flux analysis showing (**D**) OCR readouts of IL-4 treated (M(IL-4)) NIKWT BMDMs pretreated for 5 hours with 5μM B022 or vehicle (DMSO). Line graphs are combined technical (n > 20) and biological replicates with bar graphs representing each animal (n > 5 biological replicates). Data are mean +/- SEM and statistics are a Student's T-test with pvalues shown on samples close to significant **p < 0.01. (**E**) Representative western blot showing inhibition of NIK post 5μM B022 treatment. Samples were pretreated with B022 for 5 hours before treatment with 20ng/mL IL-4 for 4 hours and extracting protein with RIPA buffer.
