## Supplemental Figure 2 for "NF-κB-Inducing Kinase (NIK) Governs the Mitochondrial Respiratory Capacity, Differentiation, and Inflammatory Status of Innate Immune Cells"

### Figure S2

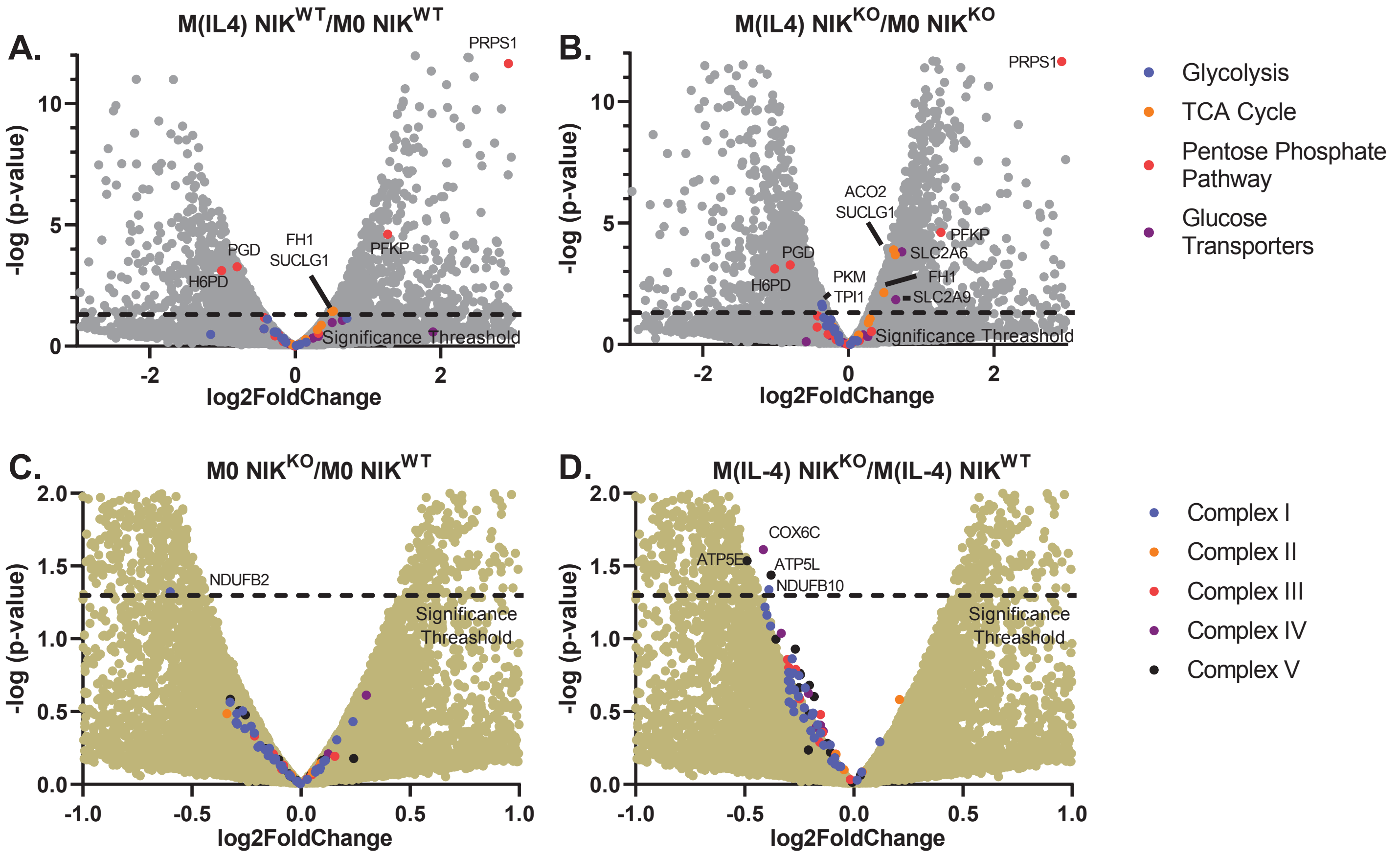


**Supplemental Figure 2: NIK$^{KO}$ macrophages have no significant transcriptional changes in metabolic genes.** RNA-sequencing heatmaps represent 2 biological replicates with a Log2Fold change post 6 hour IL-4 treatment over baseline in NIKWT (**A**) and NIKKO BMDMs (**B**) probing for changes in genes involved in glycolysis, TCA cycle, pentose phosphate pathway, and glucose transporters. RNA-sequencing heatmaps of NIKKO macrophages compared to NIKWT at baseline (**C**) or after 6 hour treatment of IL-4 (**D**) representing changes in gene expression of electron transport chain complex subunits. Statistics were calculated by DESeq2 and -log10 (p value) of 1.3 represents a p value of 0.05.
