## Supplemental Figure 3 for "NF-κB-Inducing Kinase (NIK) Governs the Mitochondrial Respiratory Capacity, Differentiation, and Inflammatory Status of Innate Immune Cells"

### Figure S3

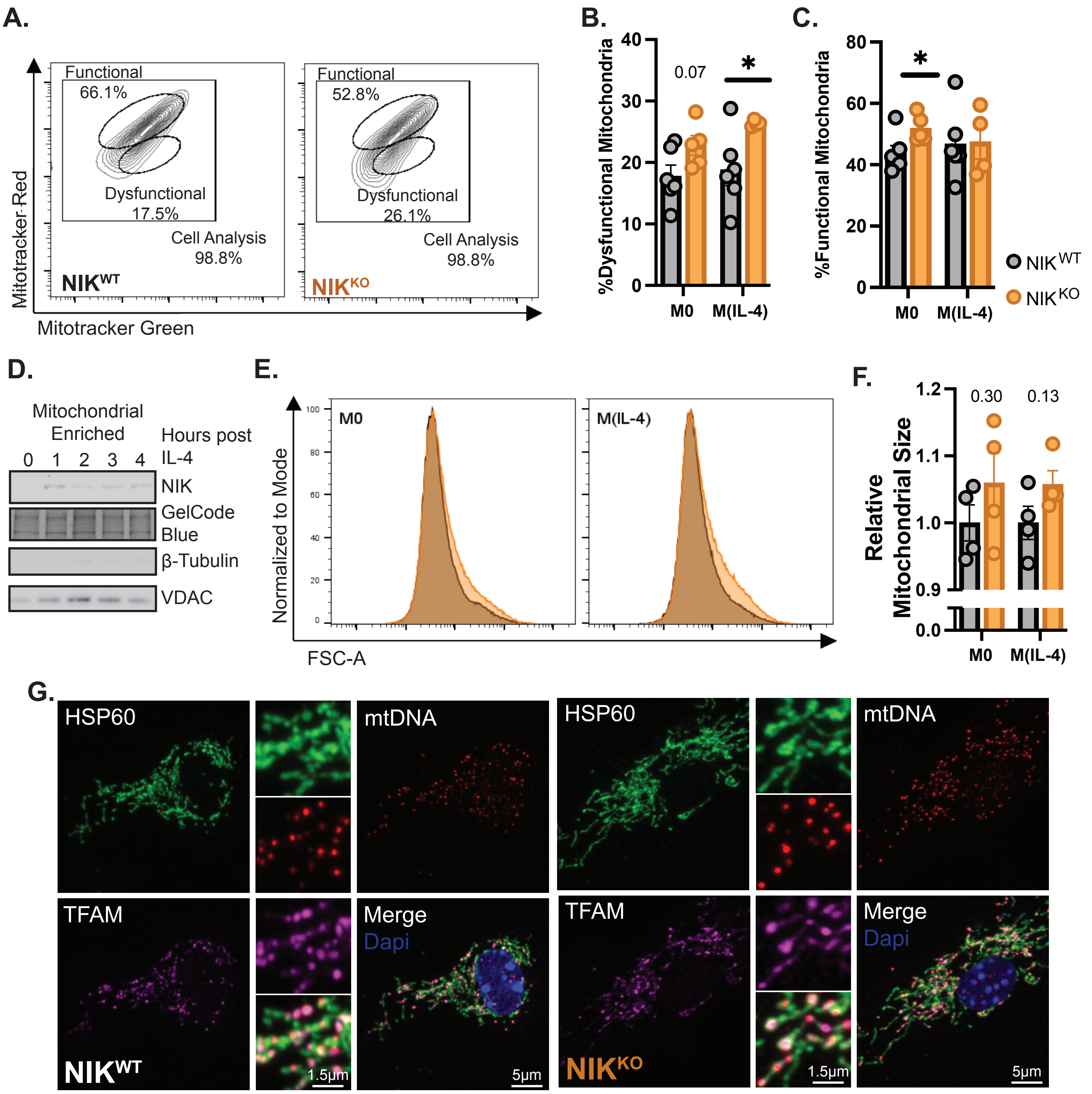


**Supplemental Figure 3: Mitochondrial homeostasis is disrupted due to a loss of NIK.** (**A**) 24 hour IL-4 stimulated BMDMs were costained with Mitotracker green and Mitotracker red to assess mitochondrial membrane potential via flow cytometry as a marker of mitochondrial functionality. Cells with low mitochondrial membrane potential were termed "dysfunctional" (**B**) and those with high mitochondrial membrane potential "functional" (**C**). Statistics are a Student's T-test. *p < 0.05 (n = 5 biological replicates). (**D**) Representative western blots from human U937 cells showing NIK localization to a mitochondrial enriched fraction post 0-4 hours of IL-4 treatment along with mitochondrial fractionation controls. (**E**) Isolated mitochondria were analyzed by flow cytometry against standard sized beads to calculate mitochondrial size with the quantification of mitochondrial size based on the standard curve of the beads (**F**) (n = 4 biological replicates). Statistics are a Student's T-test and pvalues are shown. (**G**) Representative immunofluorescent images characterizing the mitochondrial network in NIKWT and NIKKO macrophages at baseline using antibodies against HSP60 (green), mitochondrial DNA (red), TFAM (purple), and DAPI (blue).
