## Supplemental Figure 4 for "NF-κB-Inducing Kinase (NIK) Governs the Mitochondrial Respiratory Capacity, Differentiation, and Inflammatory Status of Innate Immune Cells"

### Figure S4

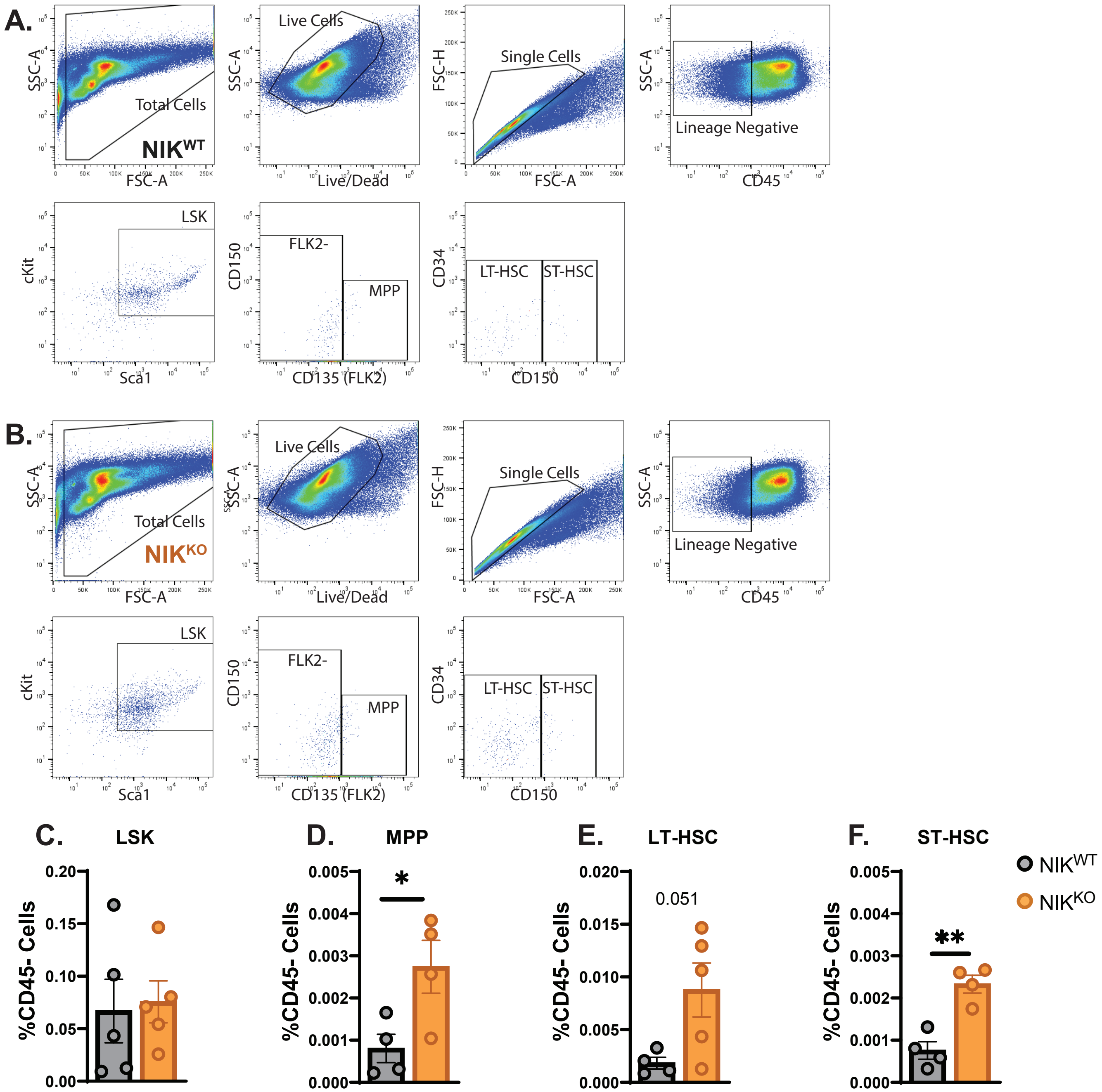


**Supplemental Figure 4: Lineage negative bone marrow cells are enriched for low proliferative cells due to a loss of NIK.** Gating scheme for stem cell analysis in NIKWT (**A**) and NIKKO (**B**) bone marrow. Quantification of the stem cell compartment probing for differences in the LSK populations (early lymphoid progenitor) (**C**), MPP (multi potent progenitor) (**D**), LT-HSC (long term hemopoietic stem cell) (**E**), and ST-HSC (short term hemopoietic stem cell) (F). Cell populations (n ≥ 4 biological replicates). Statistics are a Student's T-test. *p < 0.05, **p < 0.01.
